# Multiphoton imaging of neural structure and activity in *Drosophila* through the intact cuticle

**DOI:** 10.1101/798686

**Authors:** Max Jameson Aragon, Mengran Wang, Aaron T. Mok, Jamien Shea, Haein Kim, Nathan Barkdull, Chris Xu, Nilay Yapici

**Author notes:** These authors contributed equally to this work (Max Jameson Aragon, Mengran Wang, Aaron T. Mok, Jamien Shea). Corresponding author: Nilay Yapici.

## Abstract

We developed a multiphoton imaging method to capture neural structure and activity in behaving flies through the intact cuticles. Our measurements show that the fly head cuticle has surprisingly high transmission at wavelengths > 900 nm, and the difficulty of through-cuticle imaging is due to the air sacs and/or fat tissue underneath the head cuticle. By compressing the air sacs, we performed deep multiphoton imaging of fly brains through the intact cuticle. Our anatomical and functional imaging results show that 2- and 3-photon imaging are comparable in superficial regions such as the mushroom body, but 3-photon imaging is superior in deeper regions such as the central complex and beyond. We further demonstrated 2-photon through-cuticle functional imaging of odor-evoked calcium responses from the mushroom body γ-lobes in behaving flies short-term and long-term (12 consecutive hours). The through-cuticle imaging method developed here extends the time limits of *in vivo* imaging in flies, and opens up new ways to capture neural structure and activity from the intact fly brain.

## Introduction

Animal nervous systems across lineages have evolved to solve many of the same problems, such as foraging for food and water, finding mates to reproduce, and avoiding predators to stay alive. They navigate their environment via coordinated movements, and learn and remember the relative values of sensory stimuli around them in order to maximize their fitness and survival. At each instant in time, an animal must evaluate external sensory information and its current behavioral state to decide what to do next (Dickson, 2008; Hunt and Hayden, 2017; Lutcke et al., 2013; McFarland, 1977; Nevin, 1973; Tinbergen, 1969). A major technological challenge to revealing how the brain encodes behavioral states in real-time is that even the simplest neural computation involves interactions across the nervous system at various time scales, while our tools for assessing neural activity are restricted in time and space because of the currently available imaging sensors, methods, and preparations (Lerner et al., 2016). Optical methods remain the most established and fruitful path for revealing population dynamics in neural circuits at long time scales (ranging from minutes to hours) by providing high temporal and spatial resolution measurements (Ji et al., 2016; Luo et al., 2018; Svoboda and Yasuda, 2006).

The fly, *Drosophila melanogaster*, offers an ideal experimental system to investigate neural correlates of behavioral states and decisions because of its compact nervous system and diverse state-dependent behaviors that it executes in response to sensory stimuli (Barron et al., 2015; Dickson, 2008; Yapici et al., 2014). To understand how molecularly defined neural circuits evaluate sensory information in different behavioral states, it is critical to capture the activity of populations of neurons over long time scales as flies are changing their physiological needs (Luo et al., 2018; Simpson and Looger, 2018). These functional imaging experiments require non-invasive imaging preparations, which should allow chronic neural activity imaging for at least 12 hours. Current methods used in fly optical physiology require the fly head cuticle, trachea, and fat body to be removed by microsurgery to provide optical access to the nervous system (Grover et al., 2016; Minocci et al., 2013; Seelig et al., 2010; Sinha et al., 2013; Wang et al., 2003). These preparations are limited in imaging duration because after some time, the brain tissue starts dying due to damaged circulation resulting from the cuticle removal surgery. For example, with current imaging preparations, fly olfactory neurons show reliable Ca^2+^ responses for about four to five hours after surgery (Wang et al., 2003). A non-invasive imaging method and a preparation in which the head cuticle and underlying tissue are left intact, thereby eliminating the need for traumatic head surgery before functional imaging, is essential for advancing fly neuroscience research in the direction of chronic recordings of neural activity during ongoing behaviors. This includes being able to image the same fly brain across multiple days. In mice, multi-day imaging experiments are achieved by implanting a cranial window following removal of part of the skull (Hefendehl et al., 2012; Trachtenberg et al., 2002). Similar imaging preparations have been developed for flies (Grover et al., 2016; Huang et al., 2018; Sinha et al., 2013). However, because imaging window implantation requires a tedious surgery with low success rates and complications that occur afterwards, these methods are not commonly used. A recent development in *in vivo* multiphoton imaging is the use of long wavelength lasers in 3-photon (3P) microscopy which improves the signal-to-background ratio by several orders of magnitude compared to current 2-photon (2P) imaging methods (Horton et al., 2013; Wang et al., 2018a; Wang et al., 2018b). While 3P microscopy with 1700 nm excitation of red fluorophores and adaptive optics has shown promising results in imaging the fly brain through the cuticle (Tao et al., 2017), it is not clear if the technique is widely applicable to common blue and green fluorophores with much shorter excitation wavelengths (e.g. 1320nm).

Here, we developed a non-invasive method for imaging fly neural structure and activity through the intact head cuticle using both 2P and 3P microscopy. We first measured the ballistic and total optical transmission through the dorsal fly head cuticle and surprisingly found that the head cuticle has high transmission at the wavelengths that are used to excite green fluorophores in 2P and 3P microscopy (~920 nm and ~1320 nm, respectively). We showed that the tissue that interferes with the laser light and limits imaging through the cuticle into the brain is not the head cuticle but the air sacs and the tissue underneath the cuticle. Next, we developed a new imaging method that compresses the air sacs and allows non-invasive, through-cuticle imaging of the fly brain without any surgery. Using this imaging method, we performed deep, high spatial resolution imaging of the fly brain and determined the attenuation length for imaging through the cuticle with 2P (920 nm) and 3P (1320 nm) excitation. Our measurements showed that 2P and 3P excitation performed similarly in shallow regions of the fly brain, but 3P excitation at 1320 nm is superior for deeper brain structures. Furthermore, using 2P and 3P excitation, we imaged food odor evoked neural responses from Kenyon cells comprising the mushroom body γ-lobes using a genetically encoded Ca^2+^ indicator, GCaMP6s (Chen et al., 2013). Kenyon cells are the primary intrinsic neurons in the insect mushroom body. Diverse subtypes of Kenyon Cells (n=~2200) extend their axons along the pedunculus and in the dorsal and medial lobes (Crittenden et al., 1998; Ito et al., 1998; Strausfeld et al., 1998). These neurons receive and integrate information from heterogeneous sets of projection neurons which carry olfactory, gustatory, and visual sensory information (Owald and Waddell, 2015; Yagi et al., 2016). In our simultaneous 2P and 3P functional imaging experiments, we found no differences between 2P and 3P excitation, while recording odor evoked responses from the mushroom body γ-lobes through the cuticle. To demonstrate that our non-invasive imaging method can be used for recording neural activity in behaving flies, we used 2P excitation and captured odor evoked neural responses from mushroom body γ-lobes in flies walking on an air suspended spherical treadmill. Finally, we demonstrated long-term functional imaging by reliably capturing odor evoked neural responses from γ-lobes with 2P excitation for 12 consecutive hours. The non-invasive imaging method developed here allows multiphoton imaging of the fly brain without any microsurgery thus, opening up new ways to capture neural structure and activity from the intact fly brain at long time scales and potentially through the entire lifespan of flies.

## Results

### Fly head cuticle transmits long wavelength light with high efficiency

To develop a non-invasive imaging method using multiphoton microscopy, we first measured light transmission at different wavelengths through the fly head cuticle. Previous experiments showed that, within the wavelength range of 350 nm to 1000 nm, the relative transmission of the dorsal head cuticle of *Drosophila melanogaster* increases with increasing wavelengths (Lin et al., 2015), however, the absolute transmission, which is critical for assessing the practicality of through-cuticle imaging, was not reported. In our experiments, we quantified both the total and ballistic transmission of infrared (IR) lights through the cuticle. Dissected head cuticle samples were mounted between two glass coverslips and placed in the beam path between the laser source and the photodetector (Figure 1A). The total and ballistic transmission through the cuticle samples were measured using a custom-built system (Figure 1B). For ballistic transmission, light from a single-mode fiber was magnified and focused on the cuticle with a ~25μm spot size. Figure 1C illustrates the light path of ballistic transmission experiments. The sample stage was translated to obtain measurements at different locations on the head cuticle. Ballistic transmission through the cuticle was measured at seven different wavelengths (852 nm, 911 nm, 980 nm, 1056 nm, 1300 nm, 1552 nm, 1624 nm) that match the excitation wavelengths for typical 2P and 3P imaging. We found that for all the IR wavelengths tested, the ballistic transmission through the cuticle was high, reaching >90% at 1300 nm (Figure 1D). Since fluorescence signal within the focal volume in 2P- and 3P-microscopy is mostly generated by the ballistic photons (Dong et al., 2003; Horton et al., 2013), our results showed that ballistic photon attenuation by the fly cuticle does not limit multiphoton imaging through the intact cuticle.

**Figure 1:**
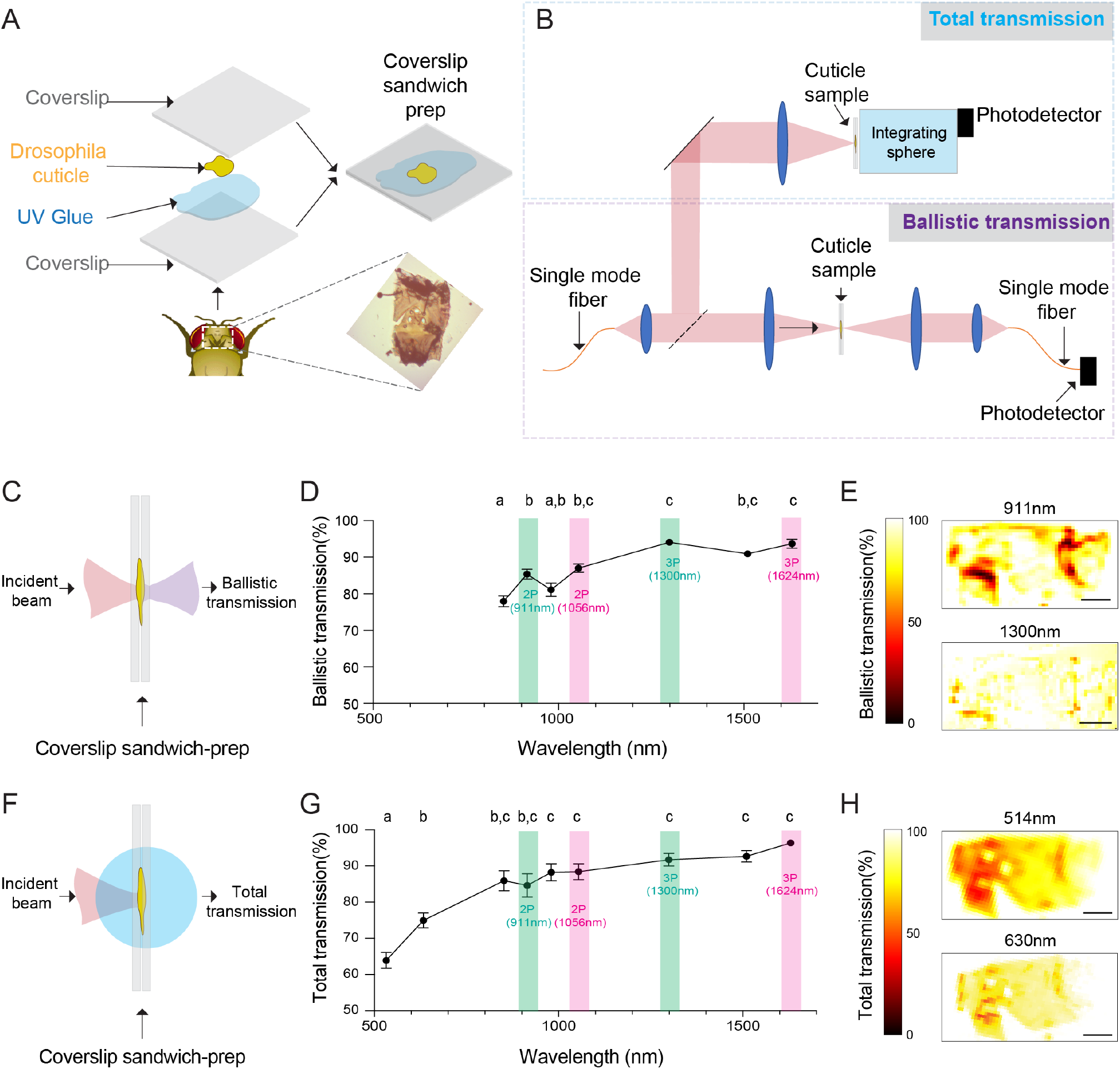
Ballistic and total optical transmission of the fly head cuticle. (**A)** Schematic of the cuticle preparation. **(B)** Schematic of the cuticle optical transmission measurement setup. (**C)** Schematic of the ballistic optical transmission through the cuticle. (**D**) Results of the ballistic optical transmission experiments at various wavelengths (n = 56 measurements at each wavelength, 5 different samples). **(E)** Spatially-resolved maps at 911 nm and 1300 nm with the percent ballistic transmission color coded. Lighter colors indicate higher transmission and darker colors indicate lower transmission. (**F)** Schematic of the total optical transmission through the cuticle. (**G**) Results of the total optical transmission experiments at various wavelengths (n = 20 measurements at each wavelength,4 different samples). (**H**) Spatially-resolved maps at 514 nm and 630 nm with the percent total transmission color coded. Lighter colors indicate higher transmission and darker colors indicate lower transmission. One-way ANOVA with post-hoc Tukey’s test. Significantly different groups are labeled with different letters in D and G. Scale bars =100μm.

To assess the absorption properties, we measured the total transmission through the head cuticle. For these measurements, laser light from a single mode fiber was magnified and focused on the cuticle sample with a ~50μm spot size. An integrating sphere was placed immediately after the cuticle to measure the total transmission. Figure 1F illustrates the light path of total transmission experiments. Total transmission through the cuticle was measured at nine different wavelengths (514 nm, 630 nm, 852 nm, 911 nm, 980 nm, 1056 nm, 1300 nm, 1552 nm, 1624 nm). The shorter wavelengths of 514 nm and 630 nm were chosen to match the typical fluorescence emission wavelengths of green and red fluorophores. Similar to the ballistic transmission experiments, we found that the total transmission generally increases with wavelength (Figure 1G), and the total transmission for both the green and red wavelengths was sufficiently high (>60%) for practical epi-fluorescence imaging using 2P or 3P excitation. We also scanned the cuticle with a motorized stage in the setup at selected wavelengths (Figure 1E, H), and these spatially-resolved transmission map confirmed that there are only a few localized regions at the periphery of the cuticle with low transmission. Our results demonstrated that absorption and scattering of long wavelength light by the *Drosophila* head cuticle is small, and non-invasive *in vivo* imaging of green (e.g., GFP and GCaMP) and red fluorophores (e.g., RFP, RCaMP) through the cuticle is possible in adult flies using 2P or 3P excitation.

### Through-cuticle structural multiphoton imaging of the fly brain

Based on our cuticle transmission results, we developed a non-invasive imaging method where we used head compression to minimize the volume of the air sacs and the tissue between the head cuticle and the brain (Figure 2A, Video 1). We imaged the fly brain in vivo through the cuticle with no head compression, semi-compression, or full-compression (Figure 2B–D). We expressed membrane-targeted GFP selectively in mushroom body Kenyon cells and imaged the fly brain through the cuticle using 2P and 3P excitation at 920 nm and 1320 nm, respectively (Figure 2E–G). Mushroom body Kenyon cells are bilaterally symmetric groups of neurons that are subdivided according to their axonal trajectories. The cell bodies of these neurons are located around the calyx of the mushroom bodies. Kenyon cell dendrites arborize in the calyx, while their axons fasciculate into anatomically distinct structures called lobes, with the dorsal lobes forming α and α′ branches, and the medial lobes containing β, β′, and γ branches (Crittenden et al., 1998; Ito et al., 1998; Zheng et al., 2018). We used transgenic flies that specifically expressed GFP in Kenyon cells forming α, β and γ lobes (Krashes et al., 2007). In non-compressed flies, the mushroom body lobes were barely visible in both 2P and 3P imaged flies. Compressing the head against the cover-glass with forceps during the curing process drastically improved image quality (Figure 2E–G). Based on our observations of the leg movements, flies behaved similarly in semi-compressed and non-compressed preps but not in full compression, and mushroom body lobes were only visible in semi-compressed preps. Thus, unless explicitly stated (e.g., for the purpose of comparisons), we used semi-compressed preps in the structural and functional imaging experiments reported in this paper.

**Figure 2.**
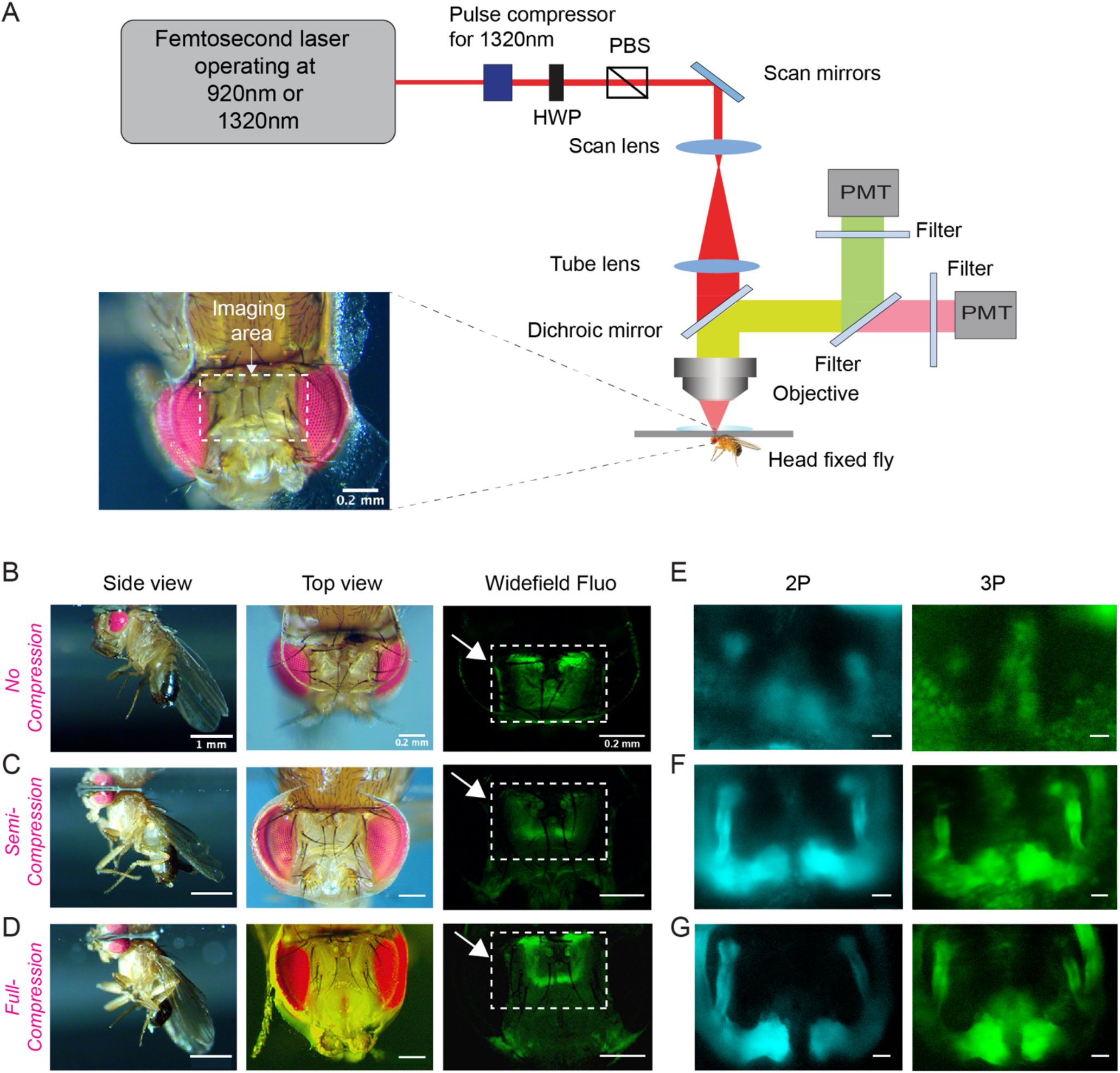
Through-cuticle imaging of the fly brain with 2P and 3P excitation. **(A)**, Schematic of the multiphoton microscope setup. Fly head is fixed to a cover slip and placed under the objective (HWP: half-wave plate, PBS: polarization beam splitter, PMT: photomultiplier tube). The imaging window on the fly head is shown in the picture (lower left). Scale bar = 200μm **(B-D)**, The head-uncompressed and head-compressed imaging preparations. The first column shows the side image of the fly that is head fixed to the cover glass (scale bar =1mm). The second and third columns show the fly head visualized under a brightfield (top view) and fluorescent dissecting microscopes (widefield-fluo), respectively. Arrows and the rectangle area in widefield-fluo column indicate the imaging window (scale bar=200μm). (**E-G)**, Cross section imaging of the mushroom body (MB) Kenyon cells expressing CD8-GFP through the head cuticle at 920 nm (2P) and 1320 nm (3P) excitation. The Z projections of 2P (cyan, left) and 3P (green, right) imaging stacks. For each imaging preparation, the same fly head is imaged with 3P excitation and 2P excitation (scale bar=20μm).

We also investigated why head compression improves image quality in intact flies. We hypothesized, head compression reduces the volume of air sacs and surrounding tissue between the cuticle and the brain, allowing better transmission of the long wavelength laser light through these structures. To test our hypothesis, we surgically removed air sacs from one side of the fly head and imaged the brain using 2P and 3P excitation without any head compression. As predicted, we were able to image the mushroom body lobes on the side where air sacs were removed but not on the side where intact air sacs were present (Figure 2- figure supplement 1A–C, Video 2). Our results demonstrated that the tissue that interferes with 2P and 3P laser light is not the cuticle itself but the air sacs and tissues that are between the head cuticle and the brain.

### Comparison of 2P and 3P excitation for deep brain imaging through the fly head cuticle

Our non-invasive imaging experiments showed that through–cuticle imaging is possible with both 2P and 3P excitation. In general, 3P excitation requires higher pulse energy at the focal plane compared to 2P excitation because of the higher-order nonlinearity. On the other hand, longer wavelength (1320 nm) used for 3P excitation can experience less attenuation while travelling in the brain tissue leading to increase tissue penetrance and imaging depth (Wang et al., 2018a). To compare the performance of 2P and 3P excitation for through-cuticle imaging, we imaged the entire brain in a fly expressing membrane-targeted GFP pan neuronally. Figure 3A shows the images from the same fly brain at different depths with 2P (920 nm) and 3P (1320 nm) excitation. At the superficial brain areas such as the mushroom bodies, 2P and 3P excitation performed similarly. As we imaged deeper in the brain, 3P excitation generated images with higher contrast compared to 2P excitation and was capable of imaging brain regions below the esophagus (~ 225 μm below the surface of head cuticle) while maintaining high spatial resolution. We further quantified the effective attenuation length (EAL) for 2P and 3P excitation, and we found EAL_920nm_= 41.7 μm, EAL_1320nm_= 59.3 μm within depth 1-100μm, and EAL_1320nm_= 91.7 μm within depth 100-180μm (Figure 3B). The third harmonic generation (THG) signal from the head cuticle and the brain trachea was also measured as a function of depth. THG signal can be used to measure the EAL (EAL_THG_) (Yildirim et al., 2019). The EAL_THG_ within the cuticle was much larger than the EAL_THG_ inside the brain, once again demonstrating the high ballistic transmission of the 1320 nm laser light through the head cuticle (Figure 3C). The full width at half maximum (FWHM) of the lateral brightness distribution at 200μm below the surface of the head cuticle was ~1.4μm for tracheal branches captured by the THG signal (Figure 3D). Similarity in the attenuation lengths measured with THG and 3P fluorescence signal indicates that the labelling of membrane-targeted GFP is uniform across the brain, validating the use of the fluorescence signal when quantifying the EALs.

To further compare the performance of 2P and 3P excitation for through-cuticle imaging in deeper brain neuropils, we imaged central complex ellipsoid body neurons. The insect central complex is a brain neuropil which processes sensory information and guides a diverse set of behavioral responses which include navigation, walking initiation, and turning direction (Pfeiffer and Homberg, 2014; Seelig and Jayaraman, 2015; Wolff et al., 2015). It is composed of anatomically distinct compartments: the protocerebral bridge, ellipsoid body, fan-shaped body, and the noduli (Wolff et al., 2015). The ellipsoid body consists of a group of neurons, the ring neurons, that extend their axons to the midline and form a ring-like structure that is composed of different layers (Pfeiffer and Homberg, 2014; Wolff et al., 2015; Xie et al., 2017). Using an ellipsoid body-specific promoter, we expressed membrane-targeted GFP in the fly brain and imaged the ring neurons with 2P and 3P excitation. Compared with 3P excitation (Figure 3E), the resolution and contrast of images taken by 2P excitation is reduced when imaging through the cuticle at this depth (Figure 3-figure supplement 1A–B). Ring neuron cell bodies, as well as the lateral triangle and the ellipsoid body layers, are clearly visible when 3P excitation was used (Figure 3E, Video 3). Using ring neuron arbors and tracheal branches, we estimated the lateral resolution of the 3P images. The FWHM of the lateral brightness distribution measured by a ring neuron’s neurite cross section was ~1.2μm for the fluorescent signal (Figure 3F) and ~0.8μm for tracheal branches captured by the THG signal (Figure 3G). Our results demonstrate that although both 2P and 3P excitation can be used to image through the cuticle for the superficial layers of the fly brain such as the mushroom body, 3P excitation outperforms 2P excitation in deeper brain regions such as the central complex. This conclusion is similar to that observed in the mouse brains (Mok et al., 2019; Wang et al., 2018a; Wang et al., 2020).

**Figure 3.**
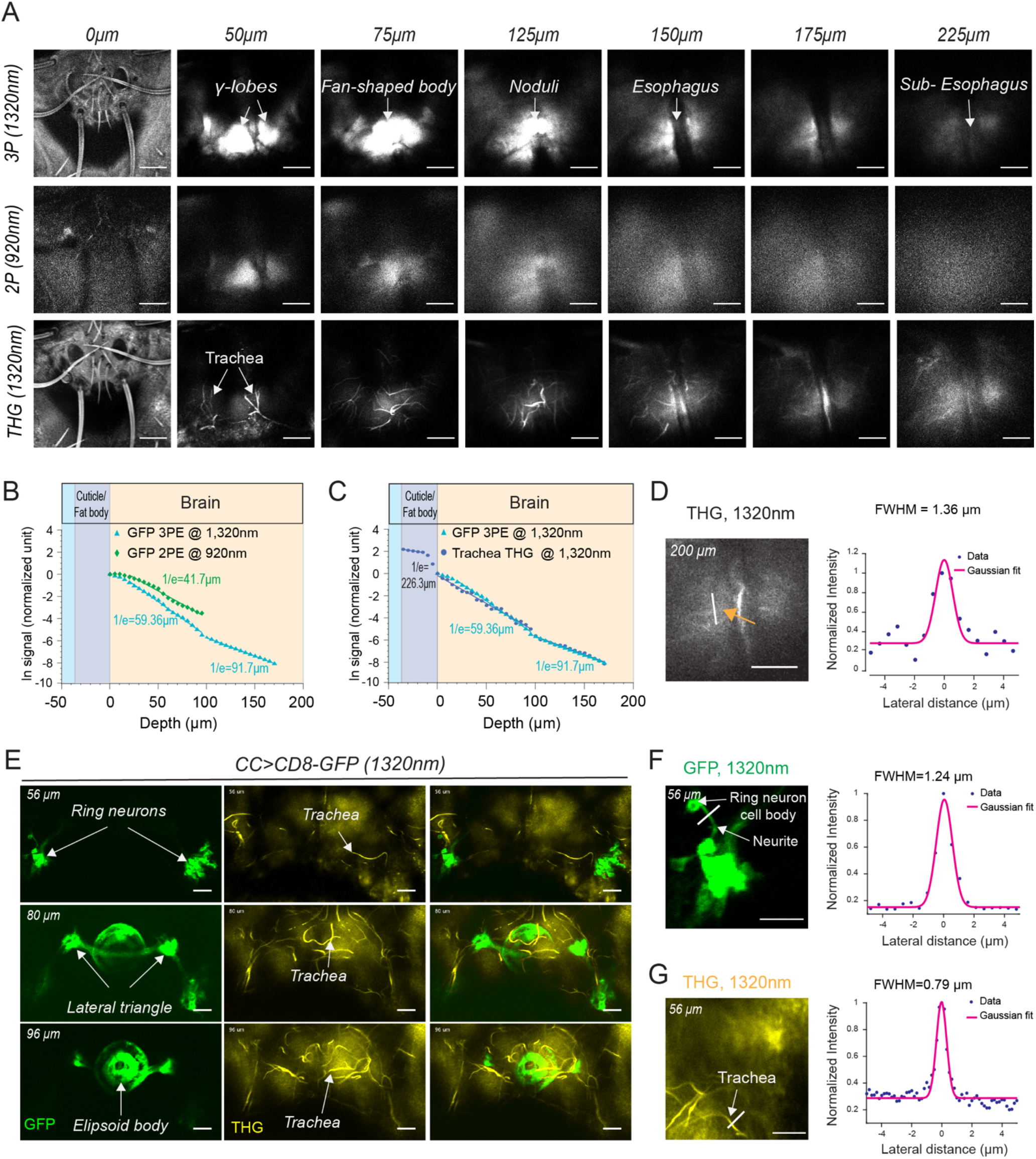
2P and 3P structural imaging of the fly brain. **(A)** Cross section images of the fly brain through the cuticle with 3P (top) and 2P (bottom) excitation at different depth. The THG images are included at the bottom. 3P excitation power is < 11mW and the repetition rate is 333 kHz. 2P excitation power is < 15mW and the repetition rate is 80 MHz. Scale bars = 50μm. (**B**) GFP signal as a function of depth for 920 nm 2P excitation and 1320 nm 3P excitation. (**C**) Comparison of the GFP signal and THG signal as a function of depth at 1320 nm. (**D)** Lateral resolution measurement in the THG image captured at 200μm depth. Lateral intensity profile measured along the white line (indicated by the orange arrow) is fitted by a Gaussian profile for the lateral resolution estimation (scale bar= 50μm). **(E)** Cross section images of the central complex (CC) ring neurons through the cuticle with 1320 nm 3P excitation (green). Third harmonic generation (THG) imaging visualizes the tracheal arbors (yellow). Arrows indicate different CC compartments that are identified (scale bars= 30μm). (**F-G**) Lateral resolution measurements in 3P images captured at 56μm depth. **(F)** The GFP fluorescence profile of CC ring neurons (green) and **(G)** the THG profile of surrounding trachea (yellow). Lateral intensity profiles measured along the white lines are fitted by Gaussian profiles for the lateral resolution estimation (scale bars=20μm).

### Simultaneous 2P and 3P imaging of odor-evoked responses from Mushroom body γ-lobes

We next compared 2P and 3P excitation to capture neural activity in the fly brain through the intact head cuticle. In these experiments, a custom odor delivery system was used where flies were head fixed and standing on a polymer ball under the microscope (Figure 4A–B). We expressed GCaMP6s selectively in Mushroom body Kenyon cells and stimulated the fly olfactory sensory organ, the antenna, with the food odor apple cider vinegar (Figure 4C–F). Using a multiphoton microscope, odor evoked Ca^2+^ responses of mushroom body γ-lobes were simultaneously captured with 2P (920 nm) and 3P (1320 nm) excitation using the temporal multiplexing technique (Ouzounov et al., 2017).

**Figure 4.**
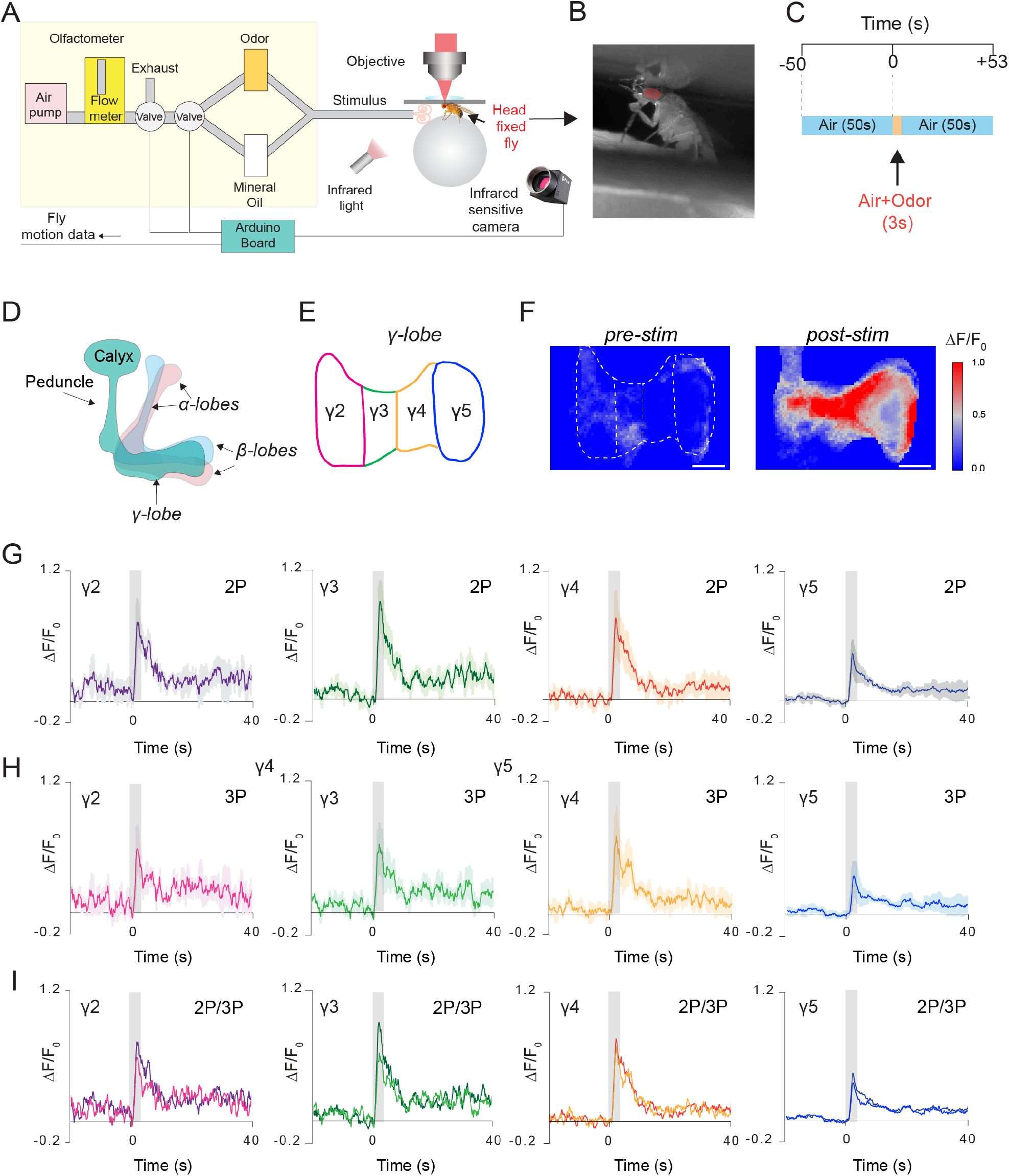
Simultaneous 2P and 3P functional imaging of short-term odor-evoked responses of the mushroom body Kenyon cells. (A) Schematic of the custom made olfactometer and the through-cuticle functional imaging setup. (**B)** Picture of the head-fixed fly on the ball under the multiphoton microscope. (**C)** Stimulus timeline. The same stimulus scheme was repeated 5 times using the same odor. (**D)** Schematic of the mushroom body anatomy indicating the locations of α, β, and γ lobes. **(E)** γ-lobes have discrete anatomical compartments (shown as γ2–γ5). **(F)** GCaMP6s is expressed in the mushroom body Kenyon cells. Normalized (ΔF/F_0_) GCaMP6s signal is shown before (left) and after (right) odor stimulus (scale bar= 20 μm). **(G)** Odor-evoked responses of Kenyon cells captured by 2P excitation at 920 nm and **(H)** 3P excitation at 1320 nm. **(I)** Comparison of the average responses captured by simultaneous 2P and 3P imaging over time (n=3 flies, 4-5 trials per fly, data are presented as mean ± SEM in (**G)** and (**H)**, grey bar indicates when stimulus is present). Average laser powers are 5 mW at 920 nm and 4 mW at 1320 nm.

A brief 3s odor stimulus triggered a robust fluorescence increase in Kenyon cell axons comprising the mushroom body γ-lobes (Figure 4F). Kenyon cell axons are innervated by distinct subsets of dopaminergic neurons, which carry contextual information to the mushroom body depending on the fly’s previous experiences. Based on dopaminergic innervation, γ-lobes can be subdivided into five anatomical compartments (Cohn et al., 2015) (Figure 4D–E). To determine whether the localized dopaminergic input along the Kenyon cell axons impacts the olfactory responses in the γ-lobes, we visualized the normalized fluorescence signal in each compartment of the γ-lobes. No significant differences were observed in neural responses to food odor stimulation across different compartments of the mushroom body or between 2P and 3P excitation of GCaMP6s (Figure 4G–I). Our data demonstrated that it is not necessary to use 3P imaging at 1320 nm for through-cuticle imaging of the mushroom body. The sample preparation technique developed here enabled conventional 2P imaging at 920 nm to capture odor evoked Ca^2+^ responses in the mushroom body through the intact cuticle with temporal resolution comparable to 2P microscopy with the open cuticle preparations (Cohn et al., 2015; Wang et al., 2003).

### Average power limit of 3P imaging in the fly brain

2P imaging has been widely used in fly neuroscience research. The recommended power level for 2P imaging of fly neural activity is below 15mW (Seelig et al., 2010). However, it is not clear, how 3P excitation impacts the fly brain. It has been shown 3P excitation can induce heating in the mouse brain at high laser powers (Wang et al., 2018a; Wang et al., 2018b; Wang et al., 2020). Therefore, we measured how heat generated by 3P excitation impacts the fly brain using HSP70 protein as a marker for cellular stress response (Lindquist, 1980; Podgorski and Ranganathan, 2016). We first tested whether HSP70 protein levels reflect heat induced stress response in the fly brain cells by placing flies in a 30°C incubator for 10 minutes and staining for HSP70 protein right after the incubation. Flies that were kept at room temperature had low levels of the HSP70 protein in a very small number of cells in the brain (Figure 4-figure supplement 1A). In contrast, placing flies in a 30°C incubator for 10 minutes caused a significant increase in the number of HSP70 positive neurons and an elevation of HSP70 protein levels across the fly brain (Figure 4-figure supplement 1B). Next, we tested whether 3P excitation can elevate HSP70 protein levels similar to heat induced stress in fly neurons. Head-fixed flies in a semi-compressed imaging prep were imaged for 10 minutes using 3P excitation at 1320 nm. Our results showed that there was no measurable heat-stress response detected by the HSP70 protein levels at the power levels < 20mW. (Figure 4-figure supplement 1C). However, increasing laser power to > 25mW caused structural abnormalities and a significant increase in HSP70 protein levels in the fly brain (Figure 4-figure supplement 2D). These results suggested that 3P non-invasive imaging of the fly brain is safe at power levels below 20mW, which is approximately 2x higher than the power we used for the deepest 3P imaging (Fig. 3A).

### 2P through-cuticle imaging captures odor evoked responses in behaving flies

To investigate how head compression impacts fly behavior and neural activity, we investigated how flies that are head-compressed but allowed to walk on a spherical treadmill respond to an odor stimulation. Using our custom behavior/imaging setup (Figure 5A), we stimulated the fly antennae that are fully exposed with food odor (apple cider vinegar), while recording neural activity from the mushroom body γ-lobes using 2P excitation (920 nm) through the head cuticle and capturing fly’s behavioral responses using a camera that is synchronized with the 2P microscope. In these experiments, a head fixed fly was continuously exposed to a low-speed air flow before and after the 3s odor stimulus with the same air flow speed, and the behavioral responses of flies were captured by tracking the spherical treadmill motion using the FicTrac software during each trial (Video 4). Because internal states impact behavioral responses to food odors (Lin et al., 2019; Sayin et al., 2019), we used flies that are 24-hour food deprived. Previous studies have demonstrated that during food odor exposure, hungry flies increase their walking speed, orient and walk towards the odor stimulus. After odor stimulation however, flies increase their turning rate which resembles local search behavior. The odor offset responses persist for multiple seconds after the odor exposure (Alvarez-Salvado et al., 2018; Sayin et al., 2019). In our experiments with semi head-compressed flies, we found that, similar to previous reports, flies increase their turning rate upon brief stimulation with food odor apple cider vinegar (Figure 5B). During these experiments, we were able to capture odor evoked neural responses from all of the mushroom body γ-lobe compartments reliably (Figure 5C).

**Figure 5.**
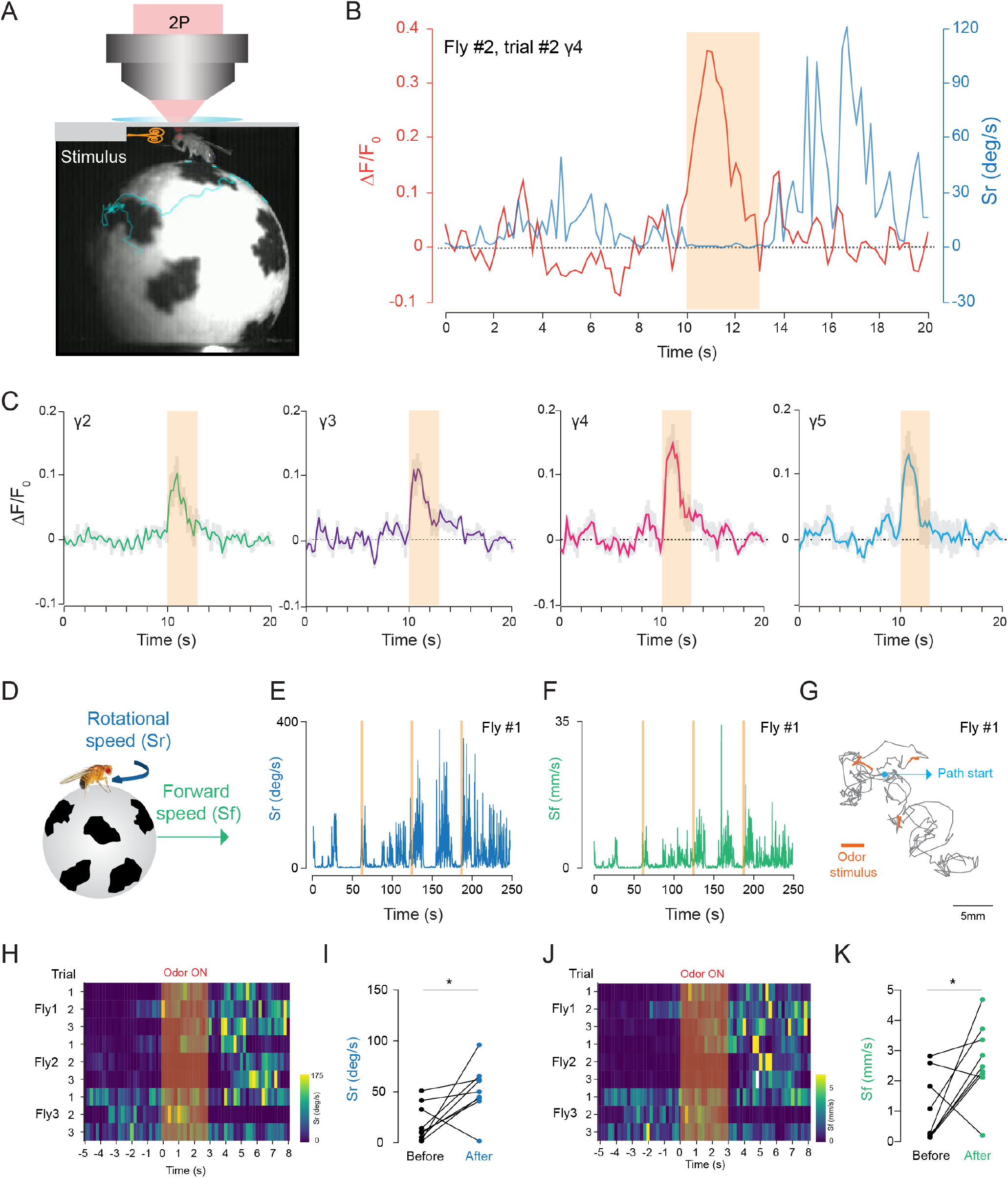
2P functional imaging of odor-evoked responses in walking flies. (A) Schematic of the custom-made odor delivery and spherical treadmill system. (**B)** Odor evoked response in mushroom body γ4 compartment is overlaid with the rotational speed (Sr) measured at the same time. **(C)** Normalized (ΔF/F_0_) GCaMP6s signal is shown during the odor stimulation experiments. Odor-evoked responses of Kenyon cells are captured by 2P excitation at 920 nm (n=3 flies, 3 trials per fly, data are presented as mean ± SEM). (**D)** Schematics showing measurements of the rotational and forward speed of flies on the spherical treadmill. (**E-G**) Representative plots for a single fly during the odor stimulation experiments showing rotational speed (Sr) (**E**), forward speed (Sf) (**F**) as a function of time, and the total calculated 2D fictive path (**G**). (**H-K**) Summary heatmap plots and statistical comparison for rotational (**H-I**) and forward speed (**J-K**) 5 seconds before and after the odor stimulation (n=3 flies, 3 trials per fly, paired-two tail t-test, p=0.0225). Average laser power at 920 nm is <10mW.

We further analyzed the odor evoked changes in fly walking behavior and showed that after the brief exposure to food odor stimulus, flies increased their forward walking speed and turning rate (Figure 5E–G). These responses lasted for multiple seconds (Figure 5H–K). Moreover, statistical analysis showed that there is a significant difference between the average forward and rotational speed values before and after the food odor exposure (Figure 5I&K). Our results are in agreement with previous studies that quantified odor induced changes in walking behavior in head fixed flies (Sayin et al., 2019). Altogether, these results indicate that head-compressed flies in our spherical treadmill setup can walk and exhibit behavioral and neural responses to odor stimulation.

### 2P through-cuticle imaging captures chronic odor evoked responses

Studying how neural circuits change activity during learning or in alternating behavioral states requires non-invasive chronic imaging methods that permit recording neural activity over long time scales. Leveraging the non-invasive nature of our preparation, we pushed the limits of functional imaging of the fly brain in response to food odor stimulation to longer time scales (12 hours). Using our custom odor delivery system, we stimulated the fly antenna with food odor (apple cider vinegar) every four hours while imaging through the head cuticle using 2P excitation (920 nm) (Figure 6A). We calculated the normalized peak fluorescent signal per fly in each γ-lobe compartment and time point as a metric representing the food odor response strength during chronic imaging (Figure 6B–E). Our analysis showed that the odor-evoked neural responses did not change with food and water deprivation in any of the γ-lobe compartments imaged (Figure 6F). During these long-term imaging experiments, we captured the fly’s behavior in parallel with the odor stimulation to assure that the fly stayed alive during long-term imaging. These results suggest that the non-invasive imaging method developed here allows recording of neural activity within an individual fly over long-time scales (12hours), which was previously not possible with commonly used open-cuticle imaging methods.

**Figure 6.**
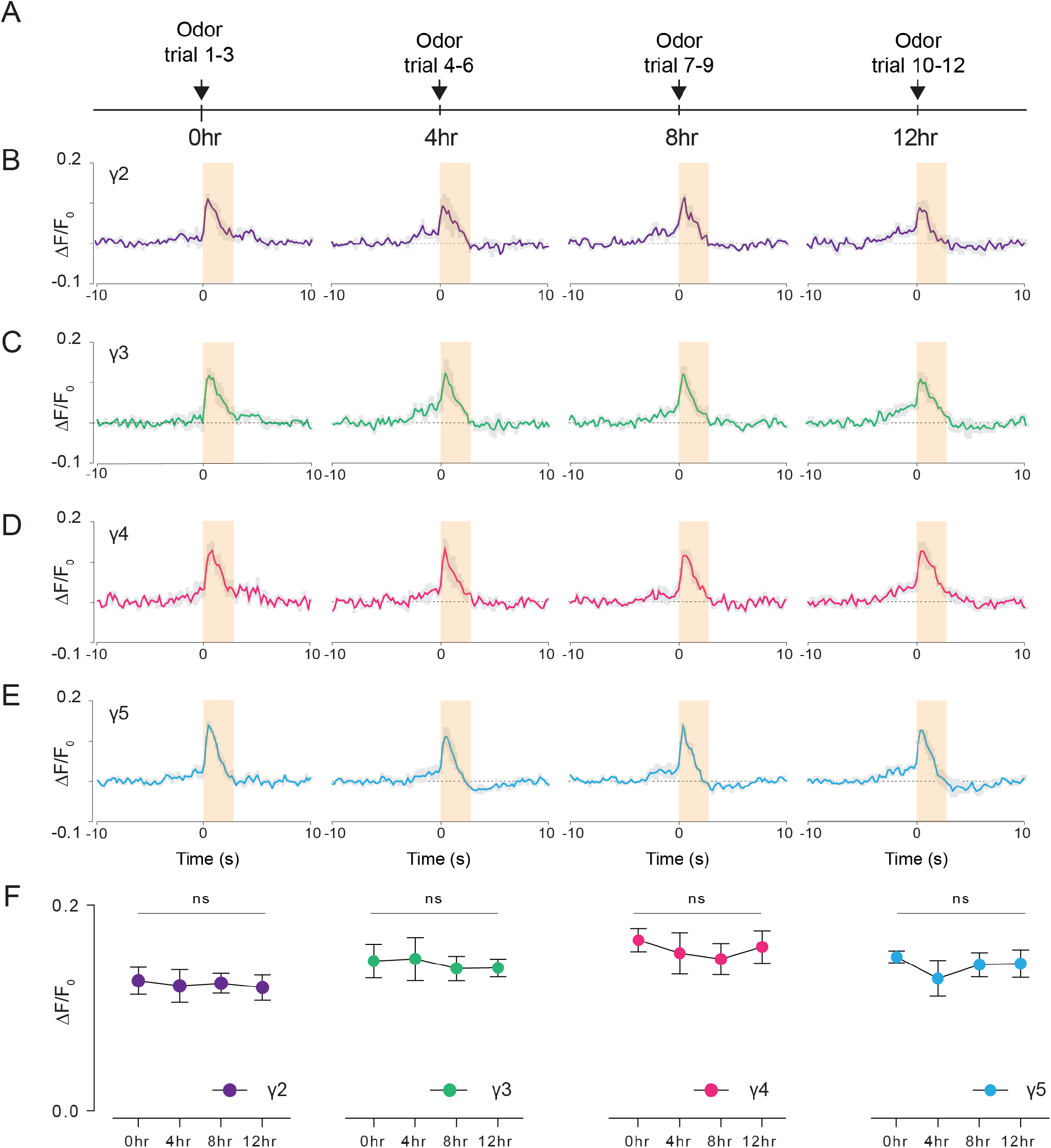
Long-term 2P imaging of odor-evoked responses of the mushroom body γ-lobes. **(A)** Stimulus timeline for long-term odor imaging. GCaMP6s fluorescence signal is captured from Kenyon cells axons innervating mushroom body γ-lobes using the semi-compressed preparation. (**B-E)** Quantification of the normalized signal (ΔF/F_0_) over time in each γ-lobe compartment. Light orange bar indicates when the odor stimulus is present. Each colored line indicates the average response of a fly over multiple trials in a given hour. The average response of 3 flies is shown. Each compartment’s response is labelled with a different color. **(F)** Quantification of the peak amplitude across different time points and lobes (ΔF_max_/F_0_) (Two-way repeated measures ANOVA. Data are presented as mean ± SEM, ns= not significant, n=3 flies, 3 trials per time point). Average laser power at 920 nm is <10mW.

## Discussion

Imaging through the fly cuticle was considered to be difficult at the wavelengths typically used for 2P (~ 920 nm) and 3P (~ 1300 nm) imaging because of concerns about cuticle absorption (Lin et al., 2015; Tao et al., 2017). By quantitatively measuring the optical properties of the fly cuticle at wavelengths that correspond to 2P and 3P fluorescence imaging, we discovered that fly cuticle transmit long wavelength light with surprisingly high efficiency (Figure 1), and it is not the absorption by the cuticle but rather the opacity of the air sacs and tissues located between the head cuticle and the brain that limit the penetration depth of multiphoton imaging (Video 2). This new understanding of the challenges for through-cuticle imaging clearly indicated that long excitation wavelength (e.g., ~ 1700 nm) is not necessary for non-invasive imaging through the fly cuticle. By compressing the fly head using a glass coverslip, we reduced the volume of the heterogenous tissues between the cuticle and the brain, which increases the transmission of laser light and therefore allows high resolution, non-invasive imaging of the fly brain through the intact cuticle (Figure 2). Careful assessments showed that such a head compression does not cause measurable differences in fly behaviors (Figure 5). Our fly preparation method enabled non-invasive 2P and 3P imaging of common fluorophores (e.g., GFP and GCaMPs) through the intact cuticle at 920 nm and 1320 nm, respectively (Figure 2–6). While we did not see noticeable differences in the recorded activity traces when performing simultaneous 2P and 3P functional imaging of the mushroom body, 3P imaging has higher contrast than 2P imaging in the deeper regions of the fly brain such as the central complex.

Investigating how physiological states, sleep, and learning change the function of neural circuits requires tracking the activity of molecularly defined sets of neurons over long time scales. These experiments require non-invasive, long term imaging methods to record neural activity *in vivo*. The non-invasive through-cuticle imaging method developed here significantly extends the time frame of current *in vivo* imaging preparations used for anatomical and functional studies in fly neuroscience. Non-invasive imaging of the fly brain will allow us to capture the activity of neural populations during changing behavioral states; facilitate decoding of neural plasticity during memory formation; and permit observation of changes in brain structures during development and aging. Our first demonstration of long-term functional imaging of the fly brain captures food odor responses from mushroom body γ-lobes for up to 12 hours continuously. Our results suggest that odor evoked Ca^2+^ responses did not change during the repeated odor stimulation. Even longer imaging time is possible by feeding flies under the microscope. We performed 2P imaging for demonstrating the possibility of long term recording of neural activity because conventional 2P microscopy has adequate penetration depth for imaging the behavioral responses within the mushroom body, and 2P microscopy is widely used by the neuroscience community. On the other hand, our deep imaging data (Figure 3A) showed that the combination of our fly preparation and 3P imaging may provide the exciting possibility of long-term imaging through the entire depth of the fly brain through the intact cuticle.

Our focus here was to develop non-invasive in vivo structural and functional imaging methods that can extend imaging quality and length for the fly brain. We anticipate that there will be a wide variety of uses for this technology in *Drosophila* neuroscience research.

## Acknowledgements

We thank Joe Fetcho, Andy Bass, David Owald, and members of the Yapici Lab for comments on the manuscript. We acknowledge Bloomington Drosophila Stock Centre (NIH P40OD018537) and the Developmental Studies Hybridoma Bank (NICHD of the NIH, University of Iowa) for reagents. We thank Li Yan McCurdy (Yale University) and Matt Einhorn (Cornell University) for help with the design and construction of the custom built olfactometer, and Nancy M. Bonini (University of Pennsylvania) for her advice on the HSP70 antibody. Research in N.Y.’s laboratory is supported by a Cornell University Nancy and Peter Meinig Family Investigator Program, a Pew Biomedical Scholar Award, the Alfred P. Sloan Foundation Award, AFAR Research Grant for Junior Faculty, NSF NeuroNex Program Grant (DBI-1707312), and NIH R35 ESI-MIRA Grant (R35GM133698-01).

## Author contributions

The authors have made the following declarations about their contributions: Conceived and supervised the study: C.X., and N.Y. Designed the experiments: M.J.A, M.W., J.S., A.M., C.X., and N.Y. Performed the experiments: J.S. and A.M. (Figure 1,5,6), M.J.A., and M.W. (Figure 2–4), H.K. (Figure 4-figure supplement 1), N.B. (Figure 1). Analyzed the data and prepared figures: M.J.A., M.W, J.S., A.M., H.K., and N.Y. Wrote the paper: N.Y. with input from C.X., M.J.A., M.W., J.S., A.M., and H.K.

## Materials and Methods

### Fly stocks

Flies were maintained on conventional cornmeal-agar-molasses medium at 23-25°C and 60-70% relative humidity, under a 12hr light: 12hr dark cycle (lights on at 9 A.M.). Fly strains and sources: *Mef2-GAL4* (Bloomington # 50742); *GMR15B07-GAL4* (Bloomington # 48678), *10XUAS-IVS-mCD8::GFP* (Bloomington # 32186); *20XUAS-IVS-GCaMP6s* (Bloomington # 42746); *GMR57C10-GAL4* (Bloomington # 39171).

### Detailed genotypes of all fly strains used in the paper are as follows

**Figure 1**

*Males, w^1118^/Y; 20XUAS-IVS-GCaMP6s; Mef2-GAL4.*

**Figure 2**

*Males, w^1118^/Y; 10XUAS-IVS-mCD8::GFP; Mef2-GAL4.*

**Figure 2-figure supplement 1**

*Males, w^1118^/Y; 10XUAS-IVS-mCD8::GFP; Mef2-GAL4.*

**Figure 3**

*Panel A-D: Males, w^1118^/Y; 10XUAS-IVS-mCD8::GFP; GMR57C10.*

*Panel E-G: Males, w^1118^/Y; 10XUAS-IVS-mCD8::GFP; GMR15B07-GAL4.*

**Figure 3-figure supplement 1**

*Males, w^1118^/Y; 10XUAS-IVS-mCD8::GFP; GMR15B07-GAL4.*

**Figure 4-6**

*Males, w^1118^/Y; 20XUAS-IVS-GCaMP6s; Mef2-GAL4.*

**Figure 4-figure supplement 1**

*Males, w^1118^/Y; 20XUAS-IVS-GCaMP6s; Mef2-GAL4.*

### Optical transmission measurements of the fly head cuticle

Drosophila cuticle was dissected from the dorsal head capsule of flies that are age and gender controlled (male, 5 days old). The dissected cuticle was sandwiched between two #1 coverslips (VWR #1 16004-094) with ~10 μL of UV curable resin (Bondic UV glue #SK8024) to avoid dehydration of the sample (Figure 1A). Measurements from each dissected cuticle was done within a day. The first several measurements were repeated at the end of all measurements to ensure that dehydration or protein degradation, which may affect the optical properties of the tissue, did not happen as the experiment progressed. The total transmission and ballistic transmission of cuticle samples were measured using a custom-built device (Figure 1B). For ballistic transmission experiments, light from a single-mode fiber was magnified and focused on the cuticle with a ~25μm spot size. We assume that collimated light passes through the sample since the Rayleigh range for a 25μm (1/e^2^) focus spot is approximately 1mm - 2mm for wavelengths between 852 nm and 1624 nm, which is much larger than the thickness of the entire coverslip sandwich-prep (<400μm). The transmitted light from the cuticle was then coupled to another single-mode fiber with identical focusing optics and detected with a power meter (S146C, Thorlabs). Such a confocal setup ensures that only the ballistic transmission is measured. The incident power is ~10mW on the cuticle for each measurement. An InGaAs camera (WiDy SWIR 640U-S, NiT) and a CMOS camera (DCC1645C, Thorlabs) were used to image the sample and incident beam to ensure that the incident light spot is always on the cuticle and to avoid the dark pigments (usually at the edge of the cuticle), ocelli, and possible cracks introduced during dissection. The ballistic transmission of the cuticle was then calculated as the power ratio between the ballistic transmissions through the cuticle (*PT^SMF^*) and the surrounding areas without the cuticle 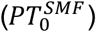, i.e., a reference transmission through areas containing only the UV curable resin, using the equation below:

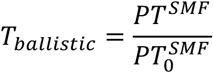

For measuring the total transmission, light from a single-mode fiber was magnified and focused on the cuticle with a ~50μm spot size. We again assume that collimated light passes through the sample since the Rayleigh range for a 50μm (1/e^2^) spot size is 5mm - 10mm for wavelengths between 532 nm and 1624 nm. An integrating sphere power meter (S146C, Thorlabs) is placed immediately after the sample to measure the total transmission. The incident power on the sample is ~10mW. The same cameras were used to visualize the light spot and the cuticle when the integrating sphere is removed. The total transmission of the cuticle was then calculated as the optical power ratio between the transmissions through the cuticle (PT^*IS*^) and the reference transmission through areas containing only the UV curable resin 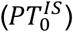, both measured by the integrating sphere (IS).

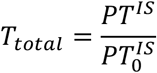

Data in Figure 1D&G are acquired by manually translating the sample orthogonal to the light path. For Figures 1E and 1H, the samples were translated with a motorized stage to acquire a spatially-resolved transmission map. We collected data from several locations for each wavelength (ballistic transmission, n=56 measurements across 5 cuticle samples; total transmission, n=20 measurements across 4 samples). We then calculated the mean and the standard error across all measurements for ballistic or total transmission for the plots shown in Figure 1D&G respectively.

### Through-cuticle *in vivo* imaging preparation

All animals used for imaging experiments were male flies with indicated genotypes kept at 25^°^C incubators and maintained on conventional cornmeal-agar-molasses medium. Flies used for chronic functional experiments were 2-7 days old, and flies used for short term functional experiments were 1-4 days old. To perform through-cuticle brain imaging, flies were first head-fixed in a 40 mm weigh dish (VWR#76299-236) with a hole made with forceps. A drop of UV curable resin (Liquid plastic welder, Bondic®) was applied to the head and thorax, which was then cured with blue light (~470 nm) and fused to a cover glass. The fly antennae are ensured to be full exposed after curing. Fly proboscises were immobilized with blue light curable resin to minimize head motion caused by muscle contractions. Video 1 explains the details of the imaging preparation. Only flies in Figure 2-figure supplement 1 went through surgery to remove the air sacs using a sharp needle.

### Multiphoton Excitation source

#### Whole brain 2P/3P imaging

The 3P excitation source is a wavelength-tunable optical parametric amplifier (NOPA, Spectra-Physics) pumped by a femtosecond laser (Spirit, Spectra-Physics) with a MOPA (Master Oscillator Power Amplifier) architecture. The center wavelength is set at 1320 nm. An SF11 prism pair (PS853, Thorlabs) is used for dispersion compensation in the system. The laser repetition rate is maintained at 333kHz. The 2P excitation source is a Ti:Sapphire laser centered at 920 nm (Chameleon, Coherent). The laser repetition rate is at 80MHz.

#### 3P imaging

The excitation source is a wavelength-tunable optical parametric amplifier (OPA, Opera-F, Coherent) pumped by a femtosecond laser (Monaco, Coherent) with a MOPA (Master Oscillator Power Amplifier) architecture. The center wavelength is set at 1320 nm. An SF10 prism pair (10SF10, Newport) is used for dispersion compensation in the system.

#### Simultaneous 2P/3P imaging and 2P imaging

The 3P excitation source is a wavelength-tunable optical parametric amplifier (NOPA, Spectra-Physics) pumped by a femtosecond laser (Spirit, Spectra-Physics) with a MOPA (Master Oscillator Power Amplifier) architecture. The center wavelength is set at 1320 nm. An SF11 prism pair (PS853, Thorlabs) is used for dispersion compensation in the system. The laser repetition rate is maintained at 400kHz. The 2P excitation source is a Ti:Sapphire laser centered at 920 nm (Tsunami, Spectra-Physics). The laser repetition rate is at 80MHz.

### Multiphoton microscopes

#### Whole brain 2P/3P imaging

It is taken with a commercial multiphoton microscope with both 2P and 3P light path (Bergamo II, Thorlabs). A high numerical aperture (NA) water immersion microscope objective (Olympus XLPLN25XWMP2, 25X, NA 1.05) is used. For GFP and THG imaging, fluorescence and THG signals are separated and directed to the detector by a 488 nm dichronic mirror (Di02-R488, Semrock) and 562nm dichronic mirror (FF562-Di03). Then the GFP and THG signals are further filtered by a 525/50 nm band-pass filter (FF03-525/50, Semrock) and 447/60 nm (FF02-447/60, Semrock) band-pass filter, respectively. The signals are finally detected by GaAsP PMTs (PMT2101, Thorlabs).

#### 3P imaging

A scan lens with 36mm focal length (LSM03-BB, Thorlabs) and a tube lens with 200mm focal length are used to conjugate the galvo mirrors to the back aperture of the objective. The same high numerical aperture (NA) water immersion microscope objective (Olympus XLPLN25XWMP2, 25X, NA 1.05) is used. Two detection channels are used to collect the fluorescence signal and the third harmonic generation (THG) signal by photomultiplier tubes (PMT) with GaAsP photocathode (H7422-40, Hamamatsu). For 3-photon imaging of GFP and GCaMP6s at 1320 nm, fluorescence signal and THG signal were filtered by a 520/60 nm band-pass filter (FF01-520/60-25, Semrock) and a 435/40 nm band-pass filter (FF02-435/40-25, Semrock), respectively. For signal sampling, the PMT current is converted to voltage and low-pass filtered (200 kHz) by a transimpedance amplifier (C7319,Hamamatsu). Analog-to-digital conversion is performed by a data acquisition card (NI PCI-6110, National Instruments). ScanImage 5.4-Vidrio Technologies (Pologruto et al., 2003) running on MATLAB (MathWorks) is used to acquire images and control a movable objective microscope (MOM, Sutter Instrument Company).

#### Simultaneous 2P/3P imaging and 2P imaging

A scan lens (SL50-3P, Thorlabs) and tube lens (TL200-3P, Thorlabs) are used to conjugate the galvo mirrors to the back aperture of the objective. The same objective (Olympus XLPLN25XWMP2, 25X, NA 1.05) is used. Fluorescence signal are detected by photomultiplier tubes (PMT) with GaAsP photocathode (H7422-40, Hamamatsu). For GFP and GCaMP6s imaging, fluorescence signal passes through a 466 nm dichronic mirror (Di02R-466) and filtered by a 520/60 nm band-pass filter (FF01-520/60-25, Semrock). For signal sampling, the PMT current is converted to voltage by a 10-MHz transimpedance amplifier (C9999, Hamamatsu). An additional 1.9-MHz low-pass filter (Minicircuts, BLP-1.9+) was used before digital sampling. Analog-to-digital conversion is performed by a data acquisition card (NI PCI-6115, National Instruments). ScanImage 3.8 -Vidrio Technologies (Pologruto et al., 2003) running on MATLAB (MathWorks) was used to acquire images.

### Simultaneous 2P and 3P imaging

Simultaneous imaging with 2P excitation and 3P excitation is achieved by temporal multiplexing of the 920 nm Ti:Sapphire laser and 1,320 nm Spirit-NOPA laser. The setup is similar to the one described in a previous study (Ouzounov et al., 2017). Briefly, two lasers were combined with a 980 nm long pass dichronic mirror (BLP01-980R-25, Semrock) and passed through the same microscope. They were spatially overlapped at the same focal position after the objective with a remote focusing module in the 2P light path. The 920 nm laser was intensity modulated with an electro-optic modulator (EOM), which was controlled by a TTL waveform generated from a signal generator (33210A, Keysight) that is triggered by the Spirit-NOPA laser. The EOM has high transmission for ~1 μs between two adjacent Spirit-NOPA laser pulses that are 2.5 μs apart. By recording the waveform from the signal generator and the PMT signal simultaneously, the 2P and 3P excited fluorescence signals can be temporally demultiplexed with postprocessing using a custom MATLAB script.

### Preparation of flies used in imaging experiments

#### Simultaneous 2P/3P functional imaging

Flies were food deprived for 16-24 hours in vials with a wet Kim wipe. Each odor stimulation trial consisted of 50 seconds of clean mineral oil, 3 seconds of undiluted apple cider vinegar stimulus, and another 50 seconds of mineral oil. Between trials, scanning was stopped for 20 seconds to minimize the risk of imaging-induced tissue stress. Five trials were performed sequentially. The images were captured at 80*160-pixel resolution and 13.6 Hz frame rate.

#### 2P functional imaging in behaving flies

Flies were head fixed using a custom 3D-printed apparatus which also holds the tube for odor delivery. In this setup, flies are allowed to walk on a spherical treadmill and turn towards the odor stimuli. The odor stimulus is located on the right side of the fly. Each odor stimulation trial consisted of 60 seconds of clean mineral oil, 3 seconds of undiluted apple cider vinegar stimulus, and another 60 seconds of mineral oil. Every change of odor triggers the acquisition software to save in a new file. The images were captured at 256*128-pixel resolution and 16.98 Hz frame rate. Three trials were performed sequentially.

#### 2P chronic functional imaging

Flies used in long term functional imaging experiments were kept on regular fly food before the first trial to assure that they were satiated. Each odor stimulation trial consisted of 60 seconds of clean mineral oil, 3 seconds of undiluted apple cider vinegar stimulus, and another 60 seconds of mineral oil. Every change of odor triggers the acquisition software to save in a new file. The images were captured at 256*128-pixel resolution and 16.98 Hz frame rate. Three trials were performed sequentially and the three-trial block was repeated every four hours. Between trial blocks, scanning was stopped, and air passing through the stimulation tube was redirected to the exhaust valve to prevent desiccation. To further prevent desiccation, flies were placed on a water-absorbing polymer bead.

### Stimulus Delivery

#### Odor delivery during 2P/3P simultaneous functional imaging

Food odor, apple cider vinegar, was delivered using a custom built olfactometer as described previously (Raccuglia et al., 2016). Clean room air was pumped (Active Aqua Air Pump, 4 Outlets, 6W, 15 L/min) into the olfactometer, and the flow rate was regulated by a mass flow controller (Aalborg GFC17). Two Arduino controlled 3-way solenoid valves (3-Way Ported Style with Circuit Board Mounts, LFAA0503110H) controlled air flow. One valve delivered the odorized airstream either to an exhaust outlet or to the main air channel, while another valve directed air flow either to the stimulus or control channel. The stimulus channel contained a 50 ml glass vial containing undiluted apple cider vinegar (volume=10 ml) (Wegmans), while the control channel contained a 50 ml glass vial containing mineral oil (volume=10 ml). Flies were placed approximately 1 cm from a clear PVC output tube (OD = 1.3 mm, ID = 0.84 mm), which passed a ~1< L/min air stream to the antennae. The odor stimulus latency was calculated before the experiments using a photo ionization detector (PID) (Aurora Scientific). We sampled odor delivery using the PID every 20 ms and found average latency to peak odor amplitude was < 100ms across 34 measurements. Flies were stimulated with air (50s), before and after the odor stimulus (odor + air, 3s). Same stimulus scheme was repeated 3 times.

#### Odor delivery during spherical treadmill and chronic imaging

An air supported spherical treadmill setup was used to record fly walking behavior during multiphoton imaging. Male flies at 5-6 days post eclosion were anesthetized on ice for about 2 minutes and mounted to a coverslip with semi-compression as described in Video 1. The cover slip was glued to a custom 3D printed holder with an internal airway in order to deliver airflow along the underside of the coverslip directly onto the antenna without interfering with the air supported ball. The air duct was positioned 90 degrees to the right of the fly about 1cm away. Clean room air was pumped (Hygger B07Y8CHXTL) into a mass flow meter set at 1L/min (Aalborg GFC17). The regulated airflow was directed through an Arduino controlled three-way solenoid pinch valve (Masterflex UX-98302-42) using 1/16” ID tubing. The valve directed the airflow either through 50ml glass vile containing 10ml of undiluted apple cider vinegar for the stimulus, or through a 50ml glass vile containing 10ml of mineral oil for the control. The latency from stimulus signal from the Arduino to odor molecules arriving at the fly’s antenna was measured using a photo-ionization detector (Aurora Scientific) prior to the experiments and found to be <200ms to peak stimulus.

### Fly behavior

The spherical treadmill was manufactured by custom milling with 6061 aluminum alloy. The treadmill has a concave surface at the end for placing the ball, which is supported by airflow. We fabricated foam balls (Last-A-Foam FR-7110, General Plastics, Burlington Way, WA USA) that are 10mm in diameter using a ball-shaped file. We drew random patterns with black ink on the foam balls to provide a high-contrast surface for the ball tracking analysis. Fly behavior was videotaped from the side to capture any movement by a CCD camera (DCC1545M, Thorlabs) equipped with a machine vision camera lens(MVL6X12Z, Thorlabs) and 950 nm long pass filter (FELH0950, Thorlabs). The acquisition frame rate for video recording was set to 8Hz under IR light illumination at 970 nm (M970L4, Thorlabs). The stimulus signal from the Arduino is captured by NI-6009 (National Instrument) using a custom script written in MATLAB 2020b (Mathworks) to synchronize with the behavior video in data analysis.

### Image processing and data analysis

#### Resolution measurements

We measured the lateral brightness distribution of small features within the fly brains using either the GFP fluorescence signal (Figure 3F) or the THG signal (Figure 3D&G). Lateral intensity profiles measured along the white lines were fitted by a Gaussian profile for the estimation of the lateral resolution.

#### Measurement of excitation light attenuation in the fly brain

The image stack was taken with 5 μm step size in depth, and the imaging power was increased with imaging depth to keep the signal level approximately constant. The signal (S) of each frame was calculated as the average of the brightest 0.25%-pixel values and then normalized by the imaging power (P) on the fly surface. The normalization is S/P^2^ and S/P^3^ for the 2P and 3P stacks, respectively. The effective attenuation length (EAL) is then derived by least-squares linear regression of the normalized fluorescence or THG signal at different imaging depth (Figure 3B&C).

#### Image processing for structural imaging

TIFF stacks containing fluorescence and THG data were processed using Fiji, an open-source platform for biological-image analysis (Schindelin et al., 2012). When necessary, stacks were registered using the TurboReg plugin.

#### Multiplex 2P and 3P functional imaging

TIFF stacks containing fluorescence data were converted to 32 bits, and pixel values were left unscaled. Lateral movement of the sample in the image series, if any, was corrected by TurboReg plug-in in ImageJ. Images acquired during the multiplexed 2P-3P imaging sessions were first median filtered with a filter radius of 10 pixels to reduce high amplitude noise. To compute ΔF/F_0_ traces, γ-lobe ROIs were first manually selected using a custom Python script. F_0_ was computed as the average of 10 frames preceding stimulus onset. The F_0_ image was then subtracted from each frame, and the resulting image was divided by F_0_. The resulting trace was then low-pass filtered by a moving mean filter with a window size of 8 frames. Data were analyzed using Python and plotted in Microsoft Excel. Peak ΔF/F_0_ was determined by the peak value within 20 frames after the odor delivery.

#### 2P functional imaging in behaving flies and chronic functional imaging

Lateral movement of the sample in the image series, if any, was corrected by TurboReg plug-in in ImageJ. A custom script written in MATLAB 2016b is used for all subsequent processing. Every 4 frames are averaged to achieve an effective frame rate of 4.25 Hz. Regions of interest (ROIs) were generated by manual segmentation of the mushroom bodies. The baselines of the activity traces (F0) for each ROIs are determined using a rolling average of 4s over the trace after excluding data points during odor stimulation. The activity traces (F) were normalized according to the formula (F − F_0_)/F_0_. The trace is finally resampled to 5Hz with spline interpolation to compile with the timing in the motion tracking trace.

#### Fly walking behavior analysis

Fly walking traces were obtained using the FicTrac (Fictive path Tracking) software as published previously (Moore et al., 2014). The ball rotation analysis was performed using the ‘sphere_map_fn’ function, which allows the use of a previously generated map of the ball to increase tracking accuracy. We post-processed the raw output generated by FicTrac. To calculate forward and rotational speeds, we used the delta rotation vectors for each axis. Then, we down-sampled raw data from 8Hz to 5Hz by averaging the values in the 200ms time-windows. The empty data points generated from down-sampling were linearly interpolated. All FicTrac analysis were done using a custom Python code.

### Immunohistochemistry for brain tissue damage assessment

To investigate laser-induced stress in the fly brain during 3-photon imaging through the intact head cuticle, 1-2-day old male homozygous MB>GCaMP6s flies were imaged continually at 1320 nm for 10 minutes. The laser was focused 70μm below the cuticle with ~20mW or ~25mW average power. Flies were head compressed to reproduce actual experimental conditions. Laser exposed brains were dissected approximately 2 hours after mounting. To test HSP70 antibody efficacy on fly brains, a positive control group of flies was first exposed to 30°C for 10 minutes in an incubator to induce HSP70 expression. Negative control group of flies was kept at room temperature. For whole mount staining, brains were dissected in phosphate-buffered saline (PBS) and incubated in 4% paraformaldehyde (PFA) in PBS for 20-30 minutes at room temperature on an orbital shaker. Tissues were washed 3-4 times over 1 hour in PBS (calcium- and magnesium-free; Lonza BioWhittaker #17-517Q) containing 0.1% Triton X-100 (PBT) at room temperature. Samples were blocked in 5% Normal Goat Serum in PBT (NGS-PBT) for 1 hour and then incubated with primary antibodies diluted in NGS-PBT for 24 hours at 4°C. Primary antibodies used were anti-GFP (Torrey Pines, TP40, rabbit polyclonal, 1:1000), anti-BRP (DSHB, nc82, mouse monoclonal, 1:20), and anti-HSP70 (Sigma, SAB5200204, rat monoclonal, 1:200). The next day, samples were washed 5-6 times over 2 hours in PBT at room temperature and incubated with secondary antibodies (Invitrogen) diluted in NGS-PBT for 24 hours at 4°C. On the third day, samples were washed 4-6 times over 2 hours in PBT at room temperature and mounted with VECTASHIELD Mounting Media (Vector Labs, Burlingame, CA, USA) using glass slides between two bridge glass coverslips. The samples were covered by a glass coverslip on top and sealed using clear nail polish. Images were acquired at 1024×1024 pixel resolution at ~1.7 μm intervals using an upright Zeiss LSM 880 laser scanning confocal microscope and Zeiss digital image processing software ZEN. Maximum projection images of Z-stacks were generated in the ZEN image-processing software. The power, pinhole size and gain values were kept the same for all imaged brains during confocal microscopy.

## Statistics

Sample sizes used in this study were based on previous literature in the field. Experimenters were not blinded in most conditions as almost all data analysis were automated and done using a standardized computer code. All statistical analysis was performed using Prism 9 Software (GraphPad, version 9.0.2). Comparisons with one variable were first analyzed using one-way ANOVA followed by Tukey’s multiple comparisons post-hoc test. Comparisons with more than one variable were first analyzed using two-way ANOVA. Comparisons with repeated measures were analyzed using a paired t-test. P values are indicated as follows: ****p < 0.0001; ***p < 0.001; **p < 0.01; and *p < 0.05. Plots labeled with different letters in a given panel are significantly different.

## Data availability

All data supporting the findings of this study is included in the paper and the supplemental files.

---

**Figure 2– figure supplement 1.**
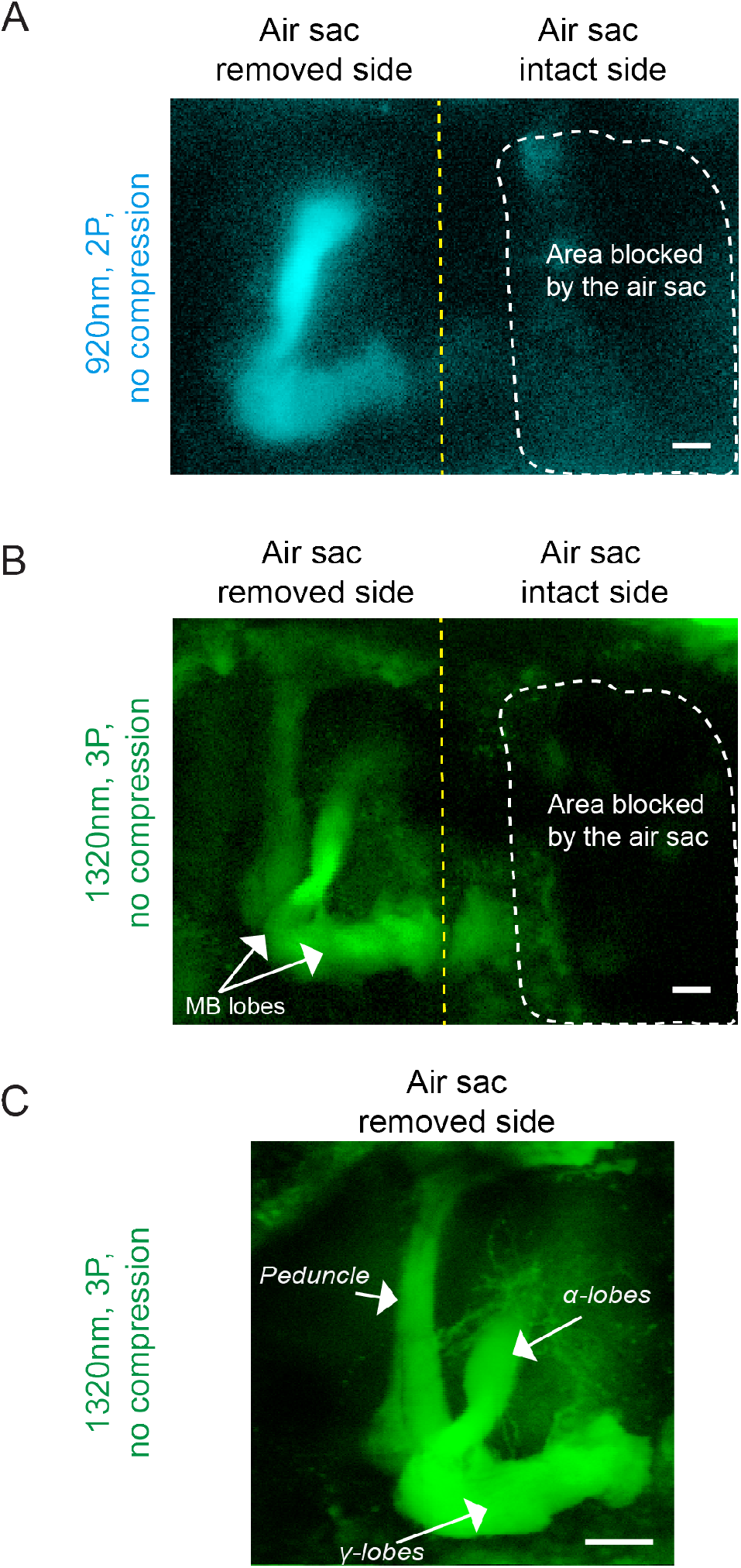
Removal of air sacs allows 2P and 3P imaging in a head-uncompressed preparation. Cross section imaging of the mushroom body (MB) Kenyon cells expressing GCaMP6s through the head cuticle at 920 nm (2P) and 1320 nm (3P) excitation after removing air sacs on one side of the head. The fly head is not compressed to the cover slip. The Z projections of **(A)** 2P (cyan) and **(B)** 3P (green) imaging stacks. For each imaging preparation, the same fly brain is imaged with 3P excitation and 2P excitation. Mushroom body structures are visible on the side where air sacs are removed. The dotted lines show the area where the air sacs block imaging. **(C)** Zoomed-in 3P image taken from the side of the brain where the air sacs are removed. Mushroom body lobes and peduncle are clearly visible without the head compression (Scale bars = 20μm).

**Figure 3– figure supplement 1.**
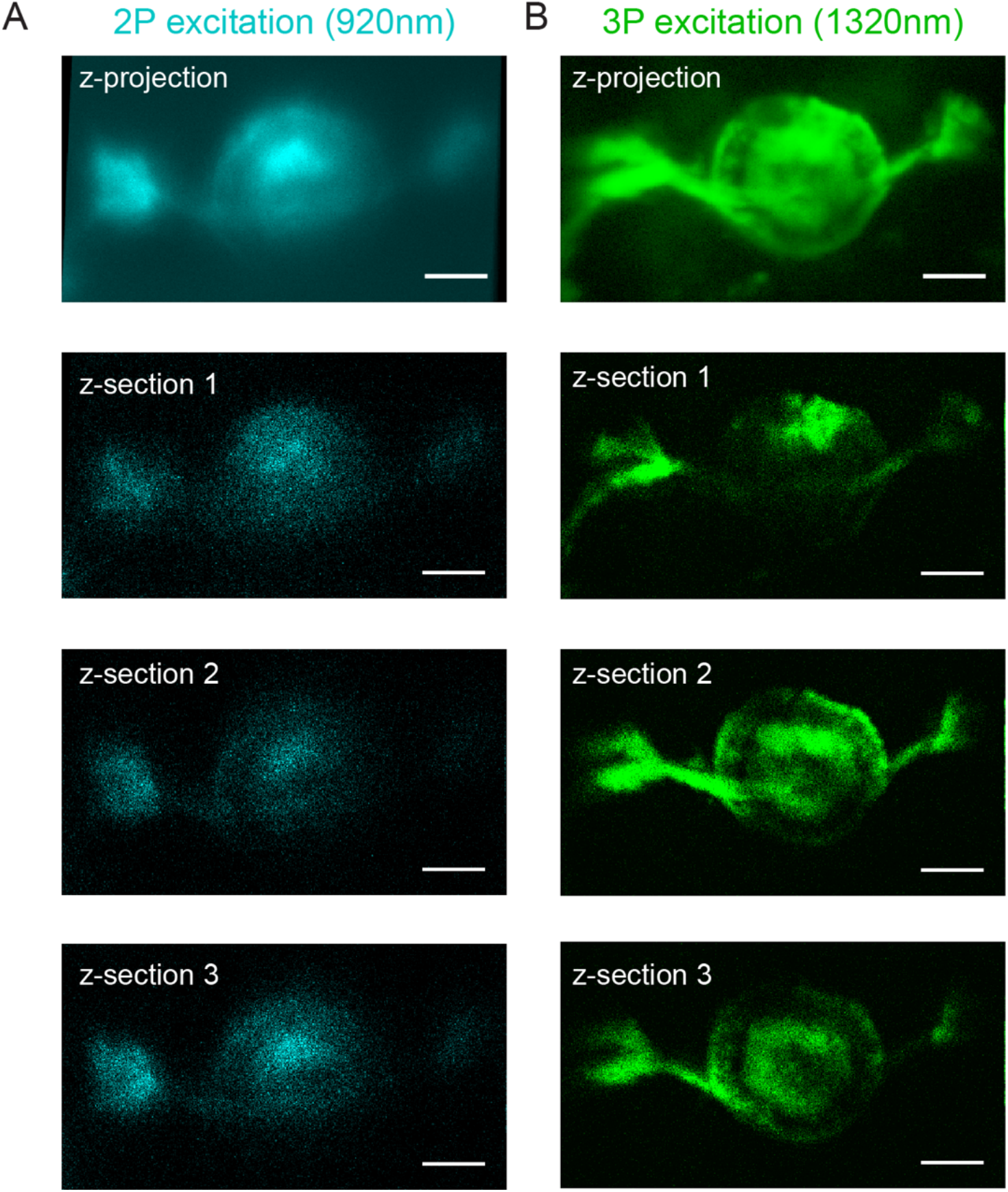
Non-invasive imaging of the central complex ring neurons with 2P and 3P excitation. Cross section imaging of the Central Complex (CC) ring neurons expressing CD8-GFP through the head cuticle at 920 nm (2P) and 1320 nm (3P) excitation. **(A-B)** The Z projection (top-most) and single optical section images at ~10 μm step of ellipsoid body processes. **(A)** 2P (cyan) and **(B)** 3P (green) images are shown. For each imaging preparation, the same fly head is imaged with both 3P and 2P excitation. Scale bars=25μm.

**Figure 4– figure supplement 1.**
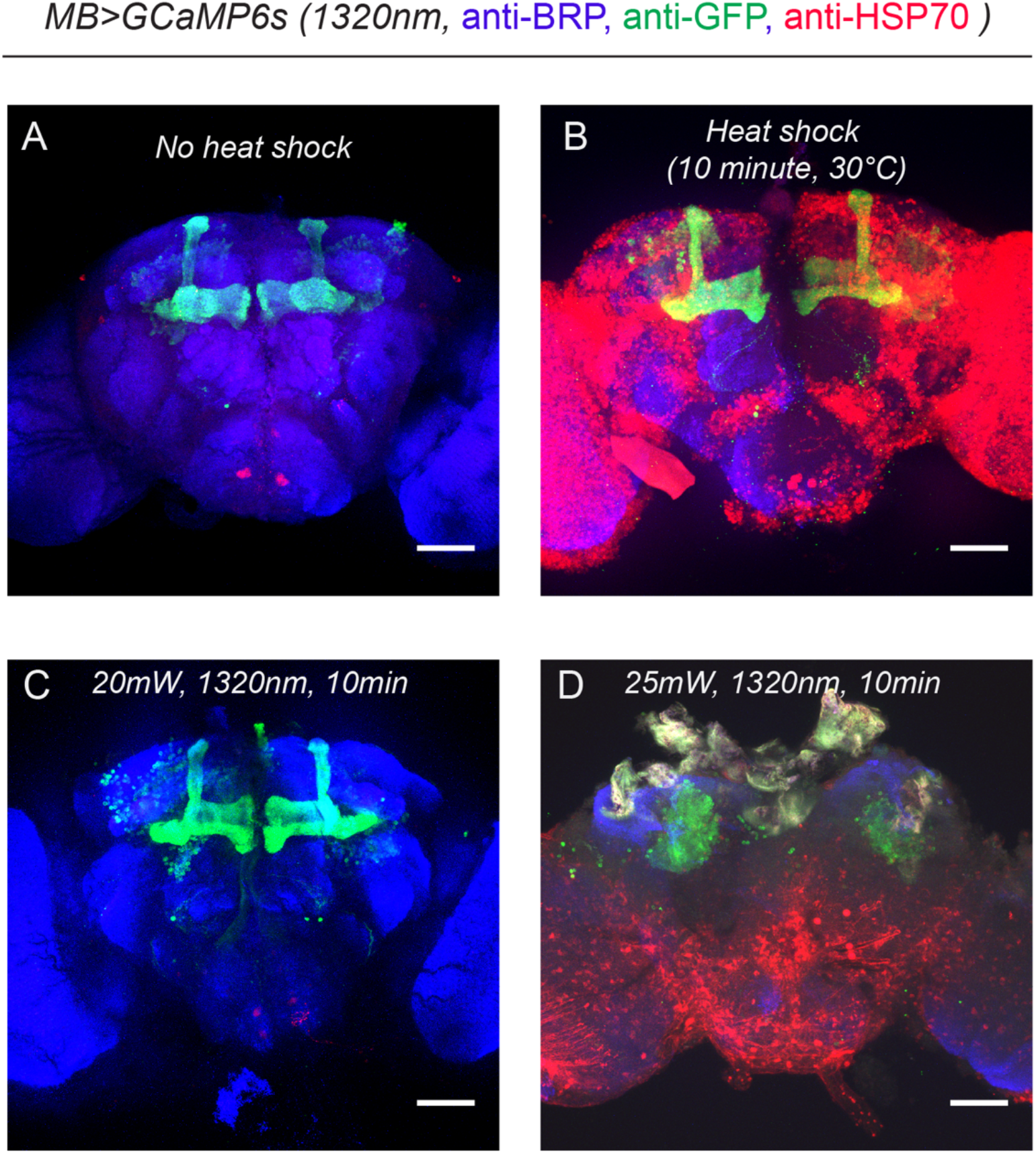
HSP70 staining of fly brains after 1320 nm 3P excitation.**(A-B)** Representative images of fly brains before and after heat shock. **(A)** Without heat shock, there is minimal HSP70 protein expressed in the fly brain. **(B)** When flies are exposed to heat shock (30°C) for 10 minutes, the HSP70 protein expression is significantly elevated across the brain. (**C-D)**, HSP70 protein expression in the fly brain after 3P excitation at 1320 nm for 10 minutes. (**C)** There is no obvious change in HSP70 protein expression when power is <20mW. (**D)** HSP70 protein expression is significantly elevated when power is >25mW. Scale bars = 50μm. The laser power is measured after the microscope objective lens using a power meter (n=2-4 flies per condition).

**Video 1: Video demonstrating how to prepare the flies for cuticle-through imaging.**

**Video 2: Z stack of the mushroom body (MB) Kenyon cells expressing GCaMP6s.** Imaging is done through the head cuticle using 1320 nm (3P) excitation after removing air sacs only on one side of the head (scale bar = 50μm, no head compression).

**Video 3: Z stack of the ellipsoid body (EB) ring neurons expressing CD8-GFP**.

Imaging through the head cuticle at 1320 nm (3P) excitation (scale bar=20μm, semi-compressed prep).

**Video 4: Neural responses of mushroom body gamma lobes captured through the intact fly head cuticle during walking and odor exposure.** Functional imaging is performed in walking flies during a food odor stimulation (apple cider vinegar) with 2P excitation at 920 nm (semi-compressed prep). Video is 5x speed up (Scale bar=30μm).

